# KIF21A influences breast cancer metastasis and survival

**DOI:** 10.1101/093047

**Authors:** Anton J. Lucanus, Victoria King, George W. Yip

## Abstract

Breast cancer pathogenesis is known to be propagated by the differential expression of a group of proteins called the Kinesin Superfamily (KIFs), which are instrumental in the intracellular transport of chromosomes along microtubules during mitosis. During mitosis, KIFs are strictly regulated through temporal synthesis so that they are only present when needed. However, their misregulation may contribute to uncontrolled cell growth due to premature sister chromatid separation, highlighting their involvement in tumorigenesis. One particular KIF, KIF21A, was recently found to promote the survival of human breast cancer cells *in vitro*. However, how KIF21A influences other cancerous phenotypes is currently unknown. This study therefore aimed to consolidate the *in vitro* role of KIF21A in breast cancer metastasis, while also analysing KIF21A expression in human breast cancer tissue to determine its prognostic value. This was achieved by silencing KIF21A in MCF-7 and MDA-MB-231 breast cancer cell lines via siRNA transfection. Migration, invasion, proliferation, and adhesion assays were then performed to measure the effects of KIF21A silencing on oncogenic behaviour. Immunohistochemistry was also conducted in 263 breast cancer tissue samples to compare KIF21A expression levels against various prognostic outcomes and clinicopathological parameters. KIF21A knockdown reduced cell migration (by 42.8% [MCF-7] and 69.7% [MDA-MB-231]) and invasion (by 72.5% [MCF-7] and 42.5% [MDA-MB-231]) in both cell lines, but had no effect on adhesion or proliferation, suggesting that KIF21A plays an important role in the early stages of breast cancer metastasis. Unexpectedly however, KIF21A was shown to negatively correlate with various pro-malignant clinicopathological parameters, including tumour size and histological grade, and high KIF21A expression predicted better breast cancer survival (hazard ratio = 0.45), suggesting that KIF21A is a tumour suppressor. The conflicting outcomes of *in vitro* and *in vivo* data may be due to the possible multi-functionality of KIF21A or study limitations, and means no definitive conclusions can be drawn about the role of KIF21A in breast cancer. This warrants further investigation, which may prove pivotal to the development of novel chemotherapeutic strategies to mediate KIF21A’s function and enhance prognostic outcomes.

## 1. INTRODUCTION

Breast cancer is the most frequently diagnosed cancer among women [1], and its metastasis often leads to death in patients. Breast cancer pathogenesis is known to be affected by the differential expression of a group of proteins called the kinesin superfamily (KIF) [2].

Kinesins were first isolated from squid tissue and identified as molecular motors for axonal transport that are ubiquitous in all eukaryotes [3]. In total, 45 kinesins have been identified in humans and other mammals [4,5]. They are sorted into 14 subfamilies based on structural differences, however all share a highly-conserved motor domain that provides motor binding to microtubules [6]. They have adenosine triphosphate (ATP) activity and microtubule-dependent motion potential, allowing movement along microtubules through coupling energy from ATP hydrolysis to force production [6].

The role of kinesins has since been specified in the transport of vesicles and organelles within cells, and chromosomes during mitosis and meiosis. During mitosis, kinesins are strictly regulated through temporal synthesis so that they are only present when needed [7]. However, misregulation of kinesins during the cell cycle may contribute to uncontrolled cell growth, highlighting their involvement in tumorigenesis. For example, *either* the depletion or overexpression of some mitotic kinesins can lead to unbalanced movement of chromosomes along microtubules during mitosis [8]. This causes a cascade of excessive spindle separation, premature sister chromatid separation, overshooting before anaphase, and finally unequal distribution of DNA and aneuploidy [9–11]. The aneuploid daughter cells may display cancerous behaviour, including increased metastatic behaviour [12].

Furthermore, Some KIFs have recently been associated with poor prognosis [13–18] and chemotherapeutic drug resistance [19–21] in breast cancer patients and cell lines, respectively. Such an involvement in mitotic deregulation, clinical outcomes and chemotherapeutic resistance in breast cancer highlights the importance of understanding the molecular mechanisms underlying the behaviour of KIFs.

One particular KIF, KIF21A, was recently found to promote lysosomal stability and the survival of human breast cancer cells *in vitro* [22]. However, how KIF21A distinctly affects other malignant phenotypes and prognostic outcomes in breast cancer is currently unknown. This study therefore aimed to (1) identify whether KIF21A is overexpressed in breast cancer tissue; (2) understand the functional behaviour of KIF21A in breast cancer cell lines and (3) evaluate the associations between KIF21A expression levels and breast cancer recurrence, survival and various clinicopathological parameters. KIF21A functional behaviour was assessed via migration, invasion, proliferation, and adhesion assays in KIF21A-silenced versus wildtype breast cancer cell lines. Immunohistochemistry staining of KIF21A in clinical IDC breast tissue microarrays was then performed to determine KIF21A expression in breast cancer tissue and to examine relationships between KIF21A expression, clinicopathological data and prognostic outcomes. We hypothesised that KIF21A shows upregulated expression in breast cancer tissue and correlates with poor prognostic outcomes, and KIF21A enhances pro-cancerous phenotypes *in vitro,* including migration, invasion, proliferation and adhesion. Identifying exactly how KIF21A is involved in breast carcinogenesis may prove pivotal to the development of chemotherapeutic targeting to mediate its function and improve prognostic outcomes.

## 2. MATERIALS AND METHODS

### 2.1 Cell Culture

MCF-7 (ATCC: HTB-22, VA, USA) and MDA-MB-231 (ATCC: HTB-2) cell lines were cultured in Dulbecco’s Modified Eagle Medium (DMEM) supplemented with 10% fetal bovine serum (FBS) (Hyclone, Logan, UT, USA) and Roswell Park Memorial Institute medium 1640 (RPMI 1640) (Hyclone) supplemented with 10% FBS, respectively. Both cell lines were incubated at 37 °C in a humidified incubator with 5% CO_2_. All cell culture was done without antibiotics.

### 2.2 Small Interfering RNA (siRNA) Transfection

Conditions used to achieve favourable silencing efficiency in both cell lines were as follows: 1 × 10^5^ (MCF-7) or 2 × 10^5^ (MDA-MB-231) cells per well were seeded with culture medium in a 6-well plate and incubated for 24 hours, which enabled the cells to reach at least 30% confluence. Transfection with Ambion Silencer Select siRNA (Ambion, Foster City, CA, USA; Table 1) was performed using Oligofectamine (Thermo Fisher Scientific, Wilmington, DE, USA) in Opti-MEM1 Reduced Serum Medium (Thermo Fisher Scientific). For MCF-7 cells, 10 μL siRNA was mixed with 175 μL Opti-MEM1 in one tube, while 5 μL Oligofectamine was mixed with 10 μL Opti-MEM1 in another tube. For MDA-MB-231 cells, 10 μL siRNA was mixed with 100 μL Opti-MEM1 in one tube, while 10 μL Oligofectamine was mixed with 100 μL Opti-MEM1 in another tube. The tubes were incubated at room temperature for five minutes, after which the contents of both were mixed and incubated again at room temperature for a further 20 minutes. The medium in each well of the 6-well plate was removed and replaced with 800 μL (MCF-7) or 780 μL (MDA-MB-231) Opti-MEM1 and 200 μL (MCF-7) or 220 μL (MDA-MB-231) of the previously-incubated siRNA-Oligofectamine mixture. The plate was then left to incubate for 8 hours at 37 °C in a humidified incubator with 5% CO_2_. 500 μL culture medium (30% FBS) was then added to each well. At 24 hours post-transfection, the medium was replaced with fresh medium (supplemented with 10% FBS). At 48 hours post-transfection, cells were harvested from each well for use in subsequent experiments. Ambion Silencer Select Scrambled siRNA was used as the negative control and Ambion Silencer Select GAPDH siRNA was used as the positive control.

**Table 1.**
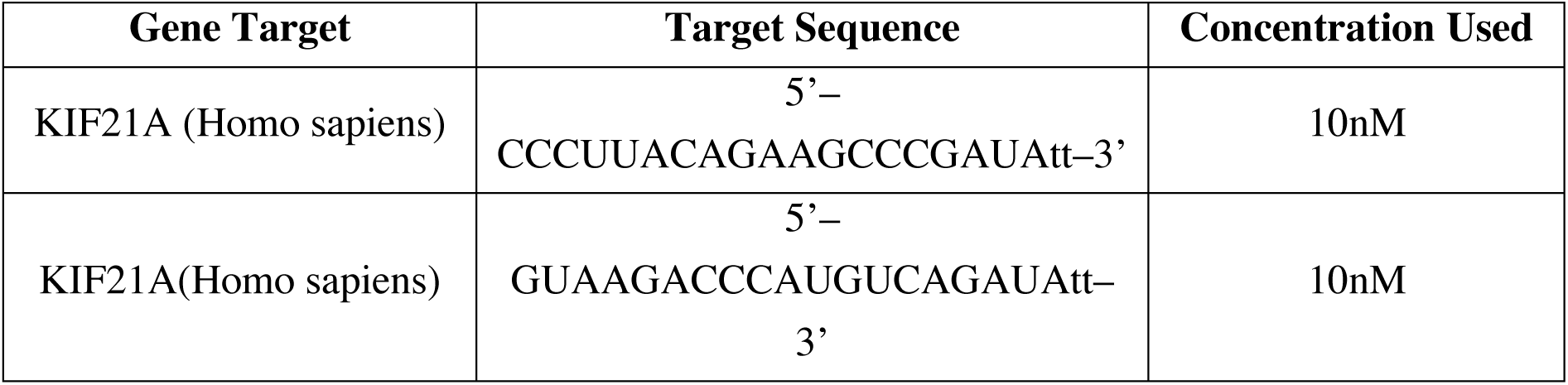
siRNAs used and their corresponding target sequences. Two different KIF21A siRNA sequences were tested to ensure reliability in results.

### 2.3 RNA Extraction and One-step qRT-PC

Total RNA was extracted using the Direct-zol RNA MiniPrep Kit (Zymo Research, Irvine, CA, USA) according to manufacturer’s instructions. RNA yield and purity were then quantified using the Nanodrop ND-100 Spectrophotometer (Thermo Fisher Scientific) according to manufacturer’s protocol. Extracted RNA was subjected to one-step qRT-PCR using the iTaq Universal SYBR Green One-Step Kit (Bio-Rad, Hercules, CA, USA), following manufacturer’s instructions, on the CFX96 Touch Real-Time PCR Detection System (Bio-Rad). The qRT-PCR primers used for this study (1st BASE, Singapore) are shown in Table 2. The program settings used for qRT-PCR were: (1) reverse transcription at 50 °C for 10 minutes; (2) activation step at 95 °C for 30 seconds; (3) 45 cycles of denaturation at 95 °C for 5 seconds and annealing at 60 °C for 30 seconds; and (4) melt curve analysis.

**Table 2.**
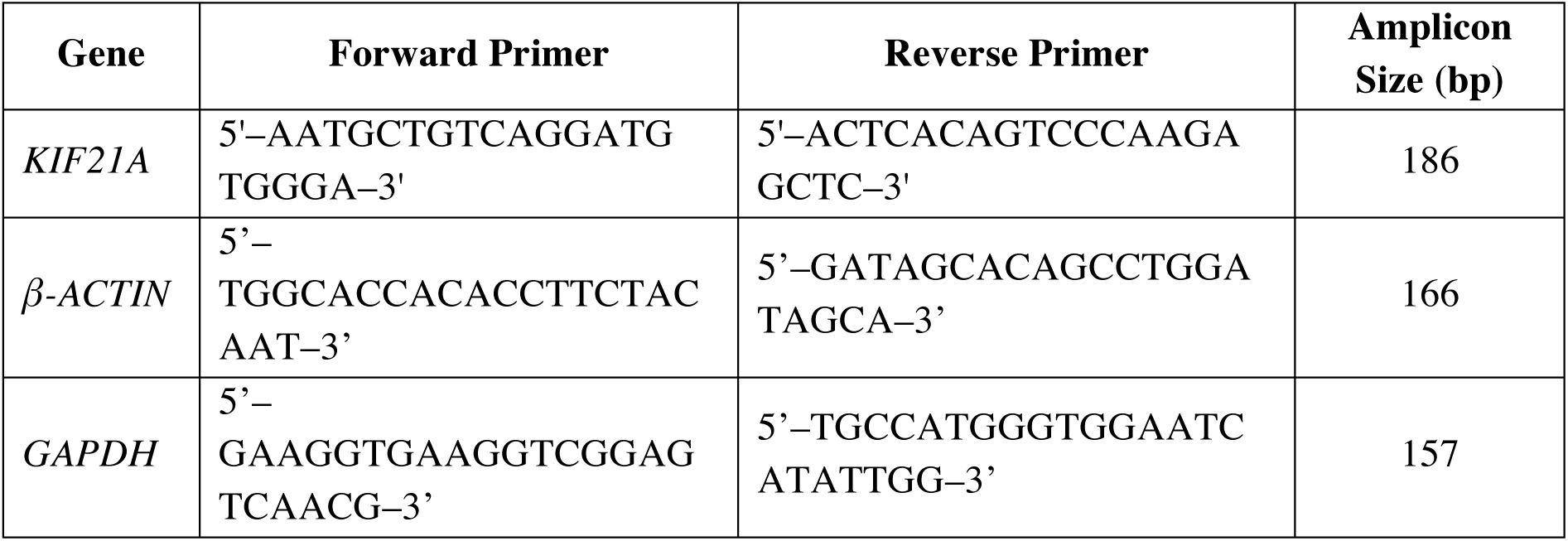
Primer sequences used during qRT-PCR.

### 2.4 Western Blotting

Protein was isolated from cells 72 hours post-transfection using M-PER Mammalian Protein Extraction Reagent (200 μL/well) (Thermo Fisher Scientific) mixed with 10 μg/mL Halt Protease Inhibitor Cocktail (2 μL/well) and EDTA (2 μL/well) (Thermo Fisher Scientific). The buffer mixture was added to cells and incubated for 5 minutes on ice. Cells were then scraped off with a cell scraper (TPP, Trasadingen, Switzerland), aspirated, and centrifuged at 16,000 *g* for 10 minutes at 4 °C. The resulting supernatant was extracted and stored at −80 °C. Following extraction, protein samples were quantified using the Bicinchoninic Acid (BCA) Protein Assay Kit (Thermo Fisher Scientific) according to manufacturer’s protocol and sodium dodecyl sulphate polyacrylamide gel electrophoresis (SDS-PAGE) was performed. Following the SDS-PAGE run, protein samples were transferred onto a polyvinylidene fluorise (PVDF) membrane (Millipore, Billerica, MA, USA) via the wet transfer method. Protein was subsequently transferred at 100 V for 1 hour at 4 °C. The membrane was then blocked with 5% BSA (Sigma-Aldrich) in 1X Tris-Buffered Saline and 1% Tween 20 (TBST) for 2 hours, and incubated at 4 °C overnight with primary antibodies (Table 4.3) that were diluted in 5% BSA. Primary antibodies were then removed and the membrane was washed with 1X TBST three times for 10 minutes each time. Secondary antibodies (Table 4.3) were then added and incubated for 1 hour at room temperature. After which, the membrane was washed with 1X TBST three times for 10 minutes each time.

**Table 3.** List of primary and secondary antibodies used in Western blotting and their dilutions. Detection of bound antibodies was performed using either the Pico or Femto Substrate Systems (Thermo Fisher Scientific). The resulting blots were scanned on X-ray films using a GS-800 Calibrated Imaging Densitometer (Bio-Rad). The optical densities of protein bands were analysed with Quantity One Version 4.1.1 software (Bio-Rad).

| Primary antibody | Catalogue number | Dilution | Secondary antibody | Catalogue number | Dilution |
| --- | --- | --- | --- | --- | --- |
| KIF21A | Aviva Systems<br>ARP33932_P050 | 1:1000 | Polyclonal<br>goat anti-rabbit HRP | Dako<br>P0448 | 1:5,000 |
| β-actin | Sigma Aldrich<br>A2228 | 1:10,000 | Anti-mouse<br>IgG HRP | GE<br>Healthcare<br>NA9310 | 1:15,000 |

### 2.5 Transwell Migration Assay

Migration assays were performed using Costar 6.5 mm Transwell chambers with 8.0 μm pore polycarbonate membrane inserts (Corning, Lowell, MA, USA). siRNA transfection was performed as per section 4.2. 48 hours post-transfection, cells were harvested, resuspended in fresh medium (10% FBS), and 5 × 10^4^ (MCF-7) or 3 × 10^4^ (MDA-MB-231) cells were seeded into the inserts. The medium outside the inserts was 600 μL fresh medium supplemented with 30% FBS. The plate was then left in an incubator for 24 hours at 37 °C. Cell suspension was then removed from inside the inserts. Migrated cells on the outer surface of the membrane of the inserts were subsequently fixed by 100% methanol and stained with 0.5% (w/v) crystal violet indicator (Sigma-Aldrich). Visualisation of the migrated cells was performed using a 10X objective lens under a Nikon SMZ1500 stereomicroscope coupled to a Nikon DXM1200F digital camera (Nikon, Minato, Tokyo, Japan). Five random field images per insert were captured for quantification.

### 2.6 Matrigel Invasion Assay

The invasion assay was performed in a similar manner to the migration assay, although the chambers used were BD BioCoat Matrigel Invasion Chambers with 8.0 μm pore size inserts (BD Biosciences, San Jose, CA, USA). siRNA transfection was performed as per section 4.2. 48 hours post-transfection, cells were harvested, resuspended in fresh medium (10% FBS), and 1 × 10^5^ (MCF-7) or 5 × 10^4^ (MDA-MB-231) cells were seeded into the inserts. The medium on the outside of the inserts was 600 μL fresh culture medium, supplemented with 30% FBS. The plate was left in an incubator for 24 hours at 37 °C. Invaded cells were then fixed, stained, imaged and quantified in an identical manner to the migration assay.

### 2.7 Serum-Starved Proliferation Assay

Proliferation assays were performed using the CellTiter 96 Aqueous One Solution Cell Proliferation Assay (Promega, Fitchburg, WI, USA). To begin, cells were serum-starved overnight prior to seeding, i.e. sub-cultured in fresh medium in a 25-cm^2^ flask without FBS. siRNA transfection was performed as per section 4.2. 72 hours post-transfection, culture medium was removed and replaced with 1.8 mL fresh medium (10% FBS) with 300 μL MTS reagent. Samples were incubated at 37 °C for 1 hour prior to measuring absorbance values once every hour for 4 hours. Formazan absorbance was detected at 490 nm by a GENios Plate Reader (Tecan, Austria).

### 2.8 Cell Adhesion Assay

96-well plates were coated overnight with either 50 μL collagen I (Corning) or 50 μL fibronectin (BD Biosciences), both at concentrations of 20 μg/mL. Collagen I and fibronectin were then removed and the wells were washed twice with 1X PBS. 100 μL 1% BSA (Sigma-Aldrich) was then added to the same wells for 1 hour at room temperature for blocking. BSA was then removed and the wells were washed twice with 1X PBS. siRNA transfection was performed as per section 4.2. 48 hours post-transfection, cells were harvested, resuspended with fresh medium and seeded into the wells at a density of 1 × 10^4^ cells for collagen I and 3 × 10^4^ cells for fibronectin wells. The 96-well plate was then incubated for 60 minutes at 37 °C for cells to adhere. Subsequently, non-adhered cells were washed off by 1X PBS. 100 μl complete medium and 20 μl MTS were then added to each well. Formazan absorbance was detected at 490 nm by a GENios Plate Reader (Tecan) once every hour for 4 hours.

### 2.9 Immunohistochemistry (IHC)

Archived, formalin-fixed, paraffin-embedded breast cancer tissue samples were received from the Department of Pathology, Singapore General Hospital (SGH). Only breast invasive ductal carcinoma (IDC) cases were used for this study. A total of 287 breast IDC cases from 1997 to 2007 were analysed, including 263 tumour tissue samples and 24 adjacent normal counterparts (all female). For the full distribution of clinicopathological data see Table 4. The collection of human tissue samples for this study received ethical approval from the Institutional Review Board, Singapore General Hospital.

**Table 4.** Distribution of Clinicopathological Data in Breast IDC Patients. Abbreviations:ER: oestrogen receptor; PR: progesterone receptor; HER2: human epidermal growth factor receptor 2.

| Clinicopathological Feature | Number of Cases (%) | Clinicopathological Feature | Number of Cases (%) |
| --- | --- | --- | --- |
| *Age* |  | *Race* |  |
| $\leq 53$ yrs old | 149 (56.7) | Chinese | 219 (83.3) |
| $> 53$ yrs old | 114 (43.3) | Non-Chinese | 44 (16.7) |
| *Tumour size* |  | *Lymphovascular invasion* |  |
| $\leq 20$ mm | 90 (34.2) | No | 204 (77.6) |
| $> 20$ mm | 168 (63.9) | Yes | 58 (22.1) |
| Not available | 5 (1.9) | Not available | 1 (0.4) |
| *Lymph node involvement* |  | *Histological grade* |  |
| No | 99 (37.6) | Grade 1 | 40 (15.2) |
| Yes | 137 (52.1) | Grade 2 | 102 (38.8) |
| Not available | 27 (10.3) | Grade 3 | 114 (43.3) |
| *ER status* |  | Not available | 7 (2.7) |
| Negative | 87 (33.1) | *PR status* |  |
| Positive | 172 (65.4) | Negative | 126 (47.9) |
| Not available | 4 (1.5) | Positive | 133 (50.6) |
| *HER2 status* |  | Not available | 4 (1.5) |
| Negative | 158 (60.1) |  |  |
| Positive | 61 (23.2) |  |  |
| Not available | 44 (16.7) |  |  |

Slides were prepared using tissue microarray (TMA) technology. The TMA slides were deparaffinised twice in Clearene (Leica Biosystems), rehydrated in a graded series of ethanol, and finally washed in distilled water. The slides were then washed in 1X Tris-buffered saline (TBS) (Bio-Rad), incubated with 3% hydrogen peroxide for 30 minutes to block endogenous peroxidase activity, and washed in 1X TBS with 1% Triton X-100 (TBS-TX) (Bio-Rad) thrice. Antigen retrieval was performed by boiling the slides at 100 °C for 20 minutes in a 500 mL solution containing 0.1 M sodium citrate, 0.1 M citric acid and distilled water. After cooling to room temperature, the slides were washed in 1X TBS-TX three times. They were then blocked for 1 hour using 1:100 dilution goat serum (Dako, Agilent Technologies, Denmark) at room temperature, followed by KIF21A primary antibody (1:100 dilution; Table 2) incubation overnight at 4 °C. The next day, the slides were washed in 1X TBS-TX, then incubated with undiluted horseradish peroxidase (HRP) conjugated secondary antibodies (polyclonal goat anti-rabbit immunoglobulins, Dako, catalogue no. K4010) for 1 hour at room temperature, and washed again in 1X TBS-TX. To visualise the staining, the slides were incubated with 3,3’-Diaminobenzidine (DAB) (Thermo Fisher Scientific) for 30 minutes, followed by counter-staining in filtered concentrated Shandon Harris haematoxylin (Thermo Fisher Scientific) for 30 seconds. The slides were finally dehydrated in a graded series of ethanol and washed twice in Clearene, followed by fixing in Permount (Thermo Fisher Scientific).

The fixed TMA slides were scanned and their staining intensities assessed using Philips Image Management System software (Philips, Amsterdam, Netherlands). KIF21A immunopositivity was scored independently by one individual followed by confirmation by a trained pathologist. The scoring criteria was based on KIF21A staining intensities with ‘0’ = no staining, ‘1+’ = weak staining intensity, ‘2+’ = moderate staining intensity, and ‘3+’ = strong staining intensity. Weighted average intensity (WAI) was then calculated, which is the average intensity of each stained cell

### 2.10 Statistical Analysis

For qRT-PCR and all functional assays, Prism 5 (GraphPad Software, La Jolla, CA, USA) was used to conduct statistical analysis. One-way ANOVAs were utilised to detect significant differences (*p* < 0.05) between wildtype cells and two groups of KIF21A-silenced cells (each group used a different siRNA sequence), followed by Tukey’s multiple comparisons post-hoc test. Immunohistochemistry results were analysed using SPSS 18.0 for Windows (SPSS Inc., Chicago, IL, USA). KIF21A immunostaining in malignant versus normal tissues were compared using the non-parametric Mann-Whitney test. Relationships between KIF21A immunoscores and nominal clinicopathological parameters were analysed using Fisher’s exact test, while relationships with ordinal parameters were analysed using Kendall’s tau-c test. For the survival analyses (recurrence and mortality), the timeline measures (in months) were: (1) OS (Overall survival) = Date of Death − Date of Diagnosis; (2) SAR (Survival after Recurrence) = Date of Death − Date of Recurrence; (3) DFS (Disease free survival) = Date of Recurrence − Date of Diagnosis. Survival analyses were performed using Kaplan-Meier analysis and the log-rank (Mantel-Cox) test, and variables that achieved statistical significance in univariate analyses (*p* < 0.05) were subsequently entered into a multivariate analysis using the Cox proportional hazards model via the backward stepwise regression method (Model A). In addition, to examine the predictive value of KIF21A expression in greater detail, the analysis included a multivariate Cox proportional hazards model that included *all* clinicopathological parameters via the enter method (Model B; Table 5.5). Statistical significance was set at *p* < 0.05 for all tests.

## 3 RESULTS

### 3.1 KIF21A Silencing in Breast Cancer Cell Lines

qRT-PCR analysis showed KIF21A was significantly silenced using two different siRNA sequences, siKIF21A-1 and siKIF21A-2 in MCF-7 cells by 39.8% (*p* < 0.01) and 73.8% (*p* < 0.001), respectively, and in MDA-MB-231 cells by 67.8% (*p* < 0.001) and 62.7% (*p* < 0.01), respectively (Figures 1A,B). Western blot analysis showed silencing via siKIF21A-1 and siKIF21A-2 translated to a reduction in protein levels in MCF-7 cells by 41.8% (*p* < 0.05) and 66.4% (*p* < 0.01), respectively, and in MDA-MB-231 cells by 58.3% (*p* < 0.05) and 69.7% (*p* < 0.01), respectively (Figures 1 C,D).

**Figure 1.**
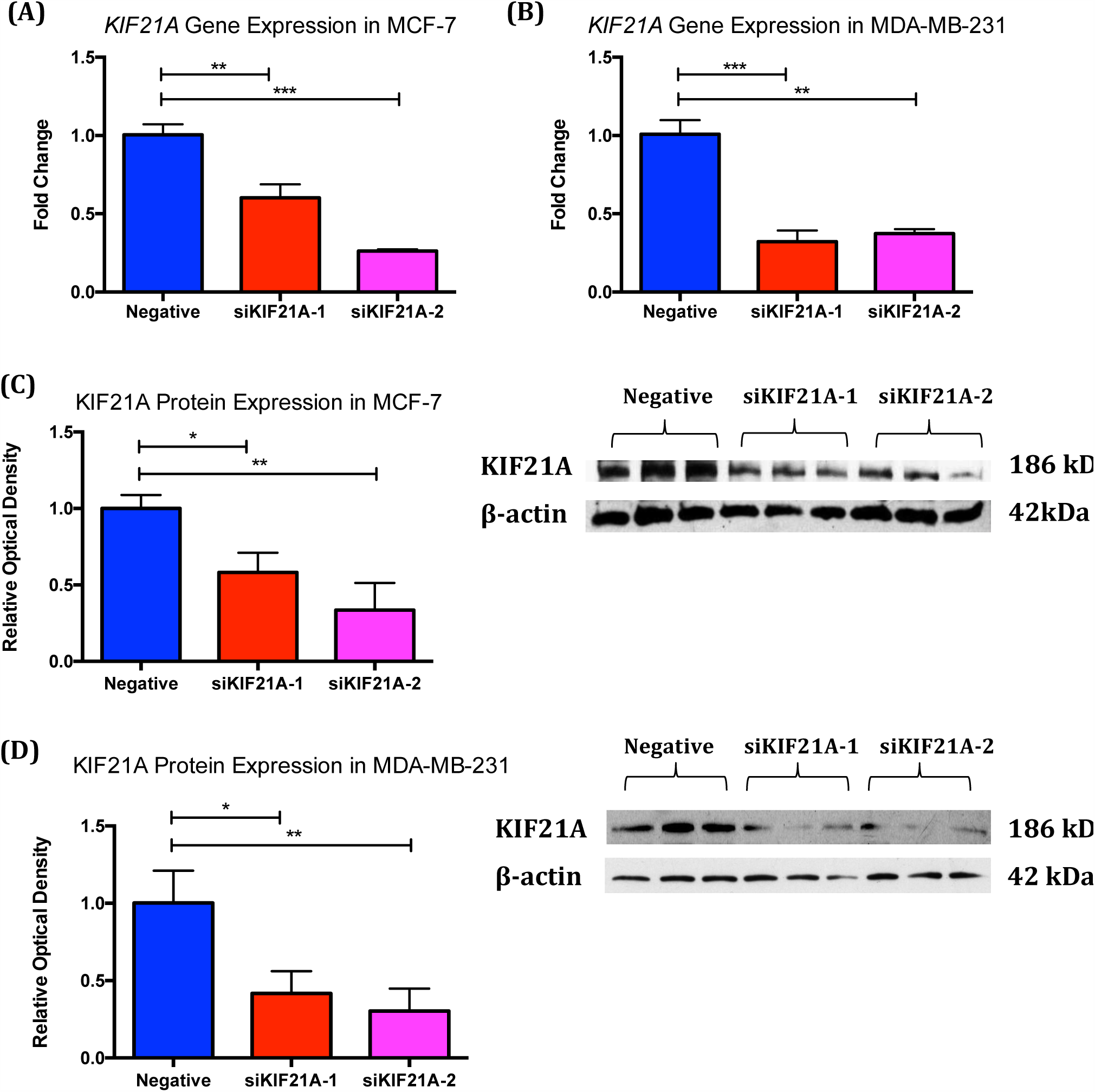
KIF21A silencing efficiencies. A,B) qRT-PCR analysis of KIF21A silencing using two siRNA sequences, siKIF21A-1 and siKIF21A-2, in MCF-7 (A) and MDA-MB-231 (B) cells showed a reduction in KIF21A mRNA expression upon silencing. C,D) Western blot protein band densitometry showed a significant reduction in the optical density of KIF21A protein bands in silenced cells for both cell lines. All protein densities from silenced cells were normalised against β-actin. For all figures: scrambled siRNA was used as the negative control. Values are mean ± SEM. n = 3 for each group, * p < 0.05, ** p < 0.01, *** p < 0.001 (One-way ANOVA with Tukey’s multiple comparisons post-hoc test).

### 3.2 Transwell Migration Assay

KIF21A silencing in MCF-7 resulted in a 42.8% (*p* < 0.01) and 39.8% (*p* < 0.01) decrease in the average number of migrated cells for siKIF21A-1 and siKIF21A-2 groups, respectively, compared to the negative control (Figures 2A,B). KIF21A silencing in MDA-MB-231 reduced cell migration by 69.7% (*p* < 0.0001) and 64.6% (*p* < 0.001) for siKIF21A-1 and siKIF21A-2, respectively, compared to the negative control (Figures 2C,D).

**Figure 2.**
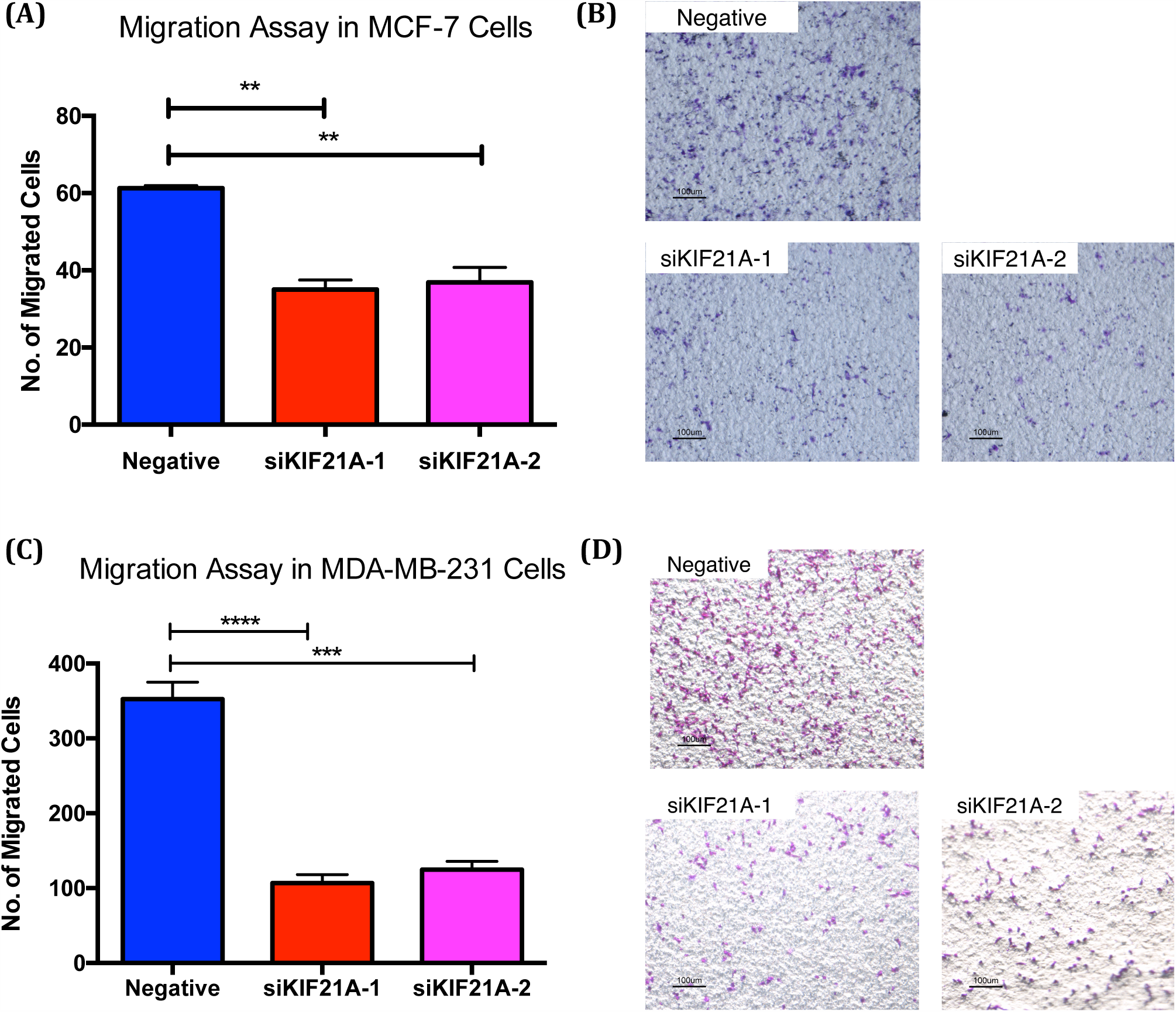
Transwell migration assays of MCF-7 and MDA-MB-231 cells following KIF21A silencing. (A,C) There was a significant decrease in migrated MCF-7 cells and MDA-MB-231 cells following KIF21A silencing for both silenced groups. (B,D) Representative photomicrographs (10x magnification) of cell migration in MCF-7 cells and MDA-MB-231 cells. For all figures: values are mean ± SEM, n = 3 for each group, ** p < 0.01, *** p < 0.001, **** p < 0.0001 (One-way ANOVA with Tukey’s multiple comparisons post-hoc test). Scale bars represent 100μm.

### 3.3 Matrigel Invasion Assay

KIF21A silencing in MCF-7 resulted in a 42.5% (*p* < 0.001) and 72.5% (*p* < 0.0001) decrease in the average number of invaded cells for siKIF21A-1 and siKIF21A-2 groups, respectively, compared to the negative control (Figures 3A,B). KIF21A silencing in MDA-MB-231 reduced cell invasion by 42.5% (*p* < 0.05) and 41.0% (*p* < 0.05) for siKIF21A-1 and siKIF21A-2, respectively, compared to the negative control (Figures 3C,D).

**Figure 3.**
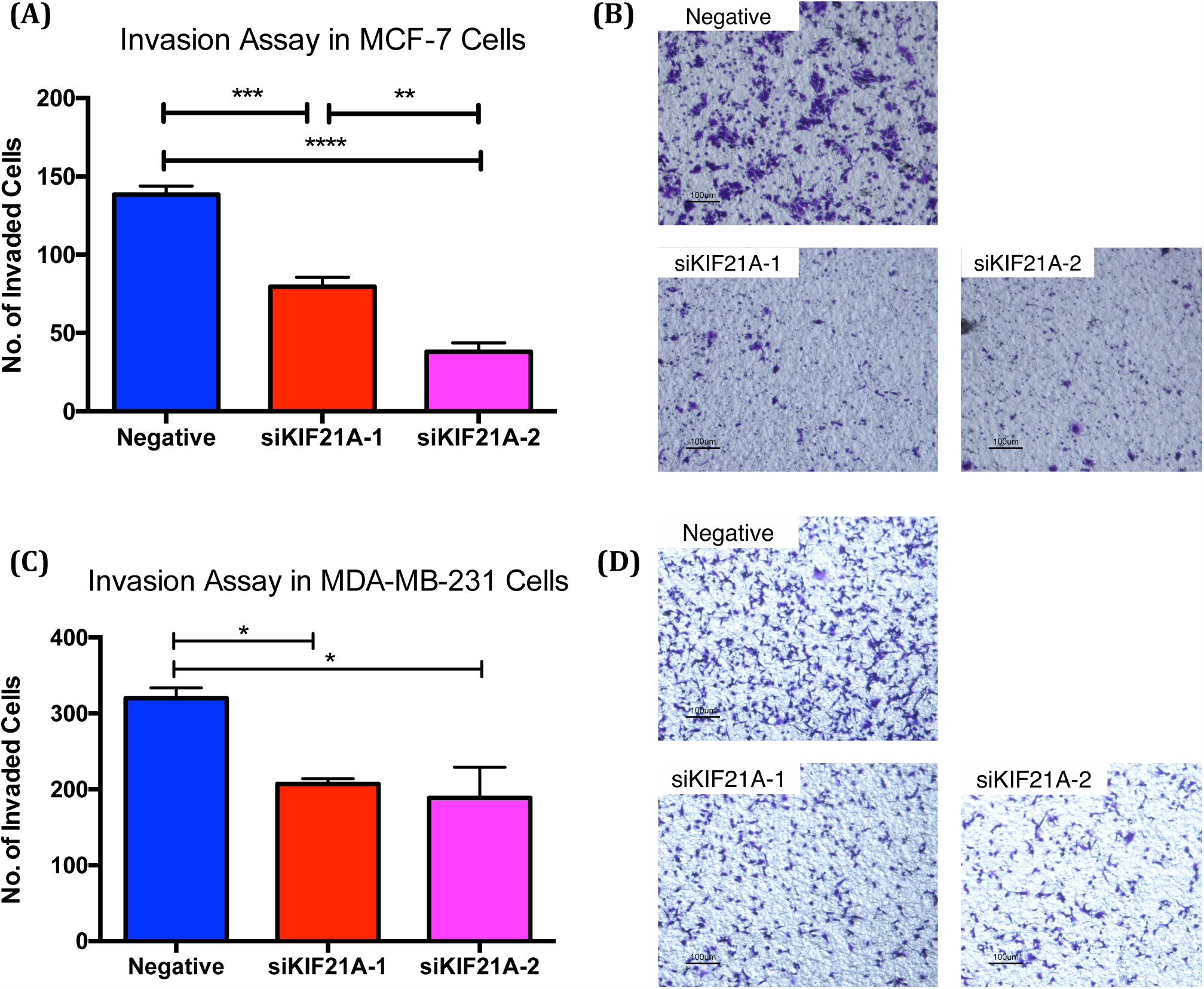
Matrigel invasion assays of MCF-7 and MDA-MB-231 cells following KIF21A silencing. (A,C) There was a significant decrease in invasive MCF-7 cells and MDA-MB-231 cells following KIF21A silencing for both silenced groups. (B,D) Representative photomicrographs (10x magnification) of cell invasion in MCF-7 cells and MDA-MB-231 cells. For all figures: values are mean ± SEM, n = 3 for each group, * p < 0.05, ** p < 0.01, *** p < 0.001, **** p < 0.0001 (One-way ANOVA with Tukey’s multiple comparisons posthoc test). Scale bars represent 100μm.

### 3.4 Cell Proliferation Assay

Analysis of cell proliferation showed no significant differences in cell proliferation between the negative and KIF21A-silenced groups for both MCF-7 and MDA-MB-231 cells (Figures 4A,B).

**Figure 4.**
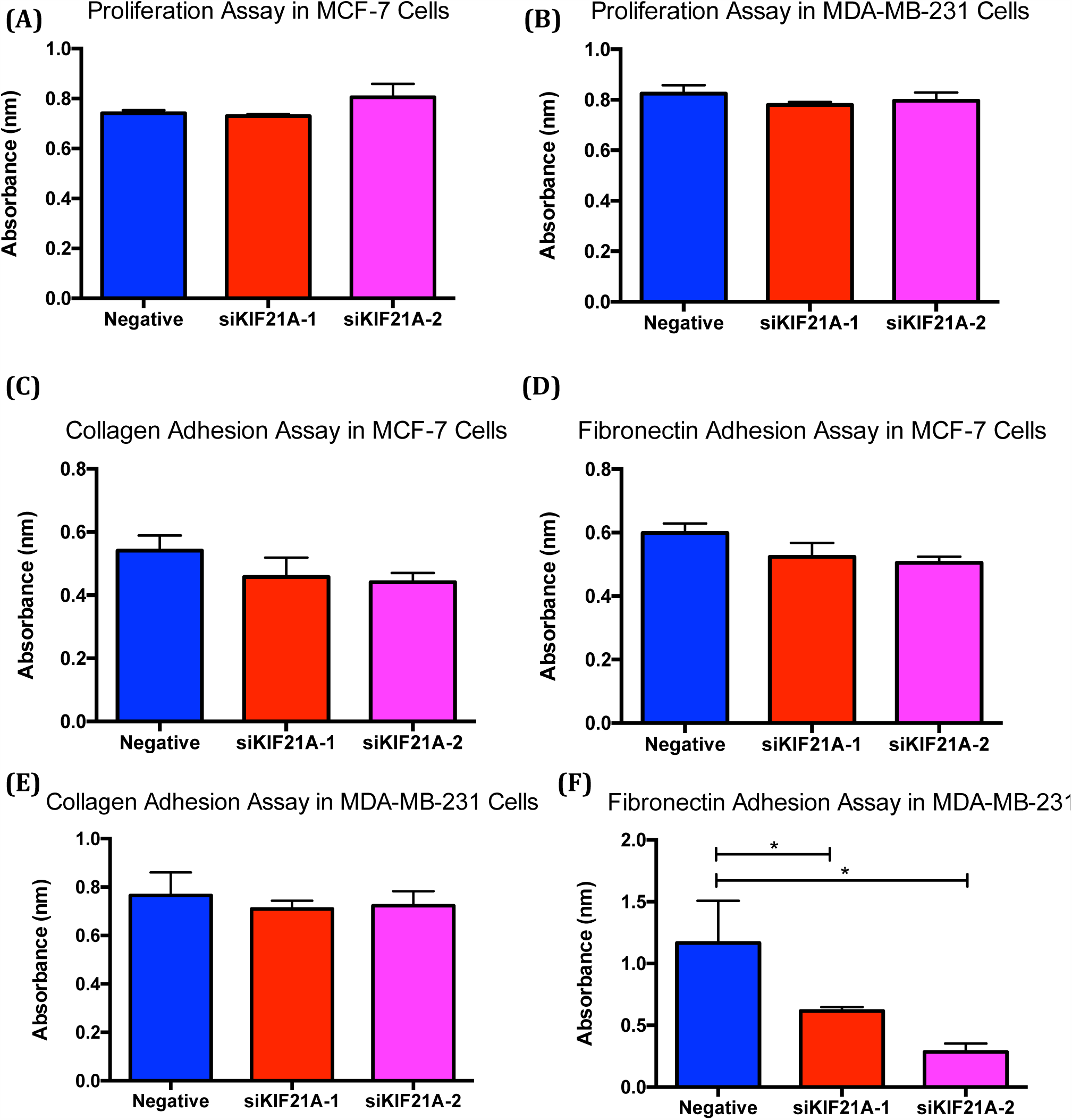
Cell proliferation and adhesion assays in MCF-7 and MDA-MB-231 cells following KIF21A silencing. (A,B) Serum-starved proliferation assay of MCF-7 and MDA-MB-231 cells following KIF21A silencing. There were no significant differences observed in cell proliferation following KIF21A silencing for both cell lines. (C,D,E,F) Adhesion assays of MCF-7 and MDA-MB-231 cells following KIF21A silencing. For all figures: absorbance (of formazan) was measured at 490 nm and indicates the relative percentage of live cells; values are mean ± SEM; n = 3 (proliferation assays) or 7 (adhesion assays) for each group, * p < 0.05 (One-way ANOVA with Tukey’s multiple comparisons post-hoc test).

### 3.5 Cell Adhesion Assay

For MCF-7 cells, analysis showed no significant differences in cell adhesion to both collagen I and fibronectin between the negative and KIF21A-silenced groups (Figures 4C,D). For MDA-MB-231 cells, KIF21A silencing also had no significant effect on cell adhesion to collagen I (Figure 4E). Interestingly, however, silenced MDA-MB-231 cells displayed reduced adhesion to fibronectin by 47.2% (*p* < 0.05) and 54.0% (*p* < 0.05) for siKIF21A-1 and siKIF21A-2, respectively (Figure 4F).

### 3.6 Comparison of KIF21A Staining Between Normal and Tumorous Breast IDC Tissues

KIF21A staining was predominately observed in the nucleus of both normal and tumourous breast epithelial cells, while cytoplasmic staining was either absent or minimal (Figure 5A,B,C). Analysis showed no significant differences in KIF21A nuclear staining between tumour and normal samples (Figure 5D).

**Figure 5.**
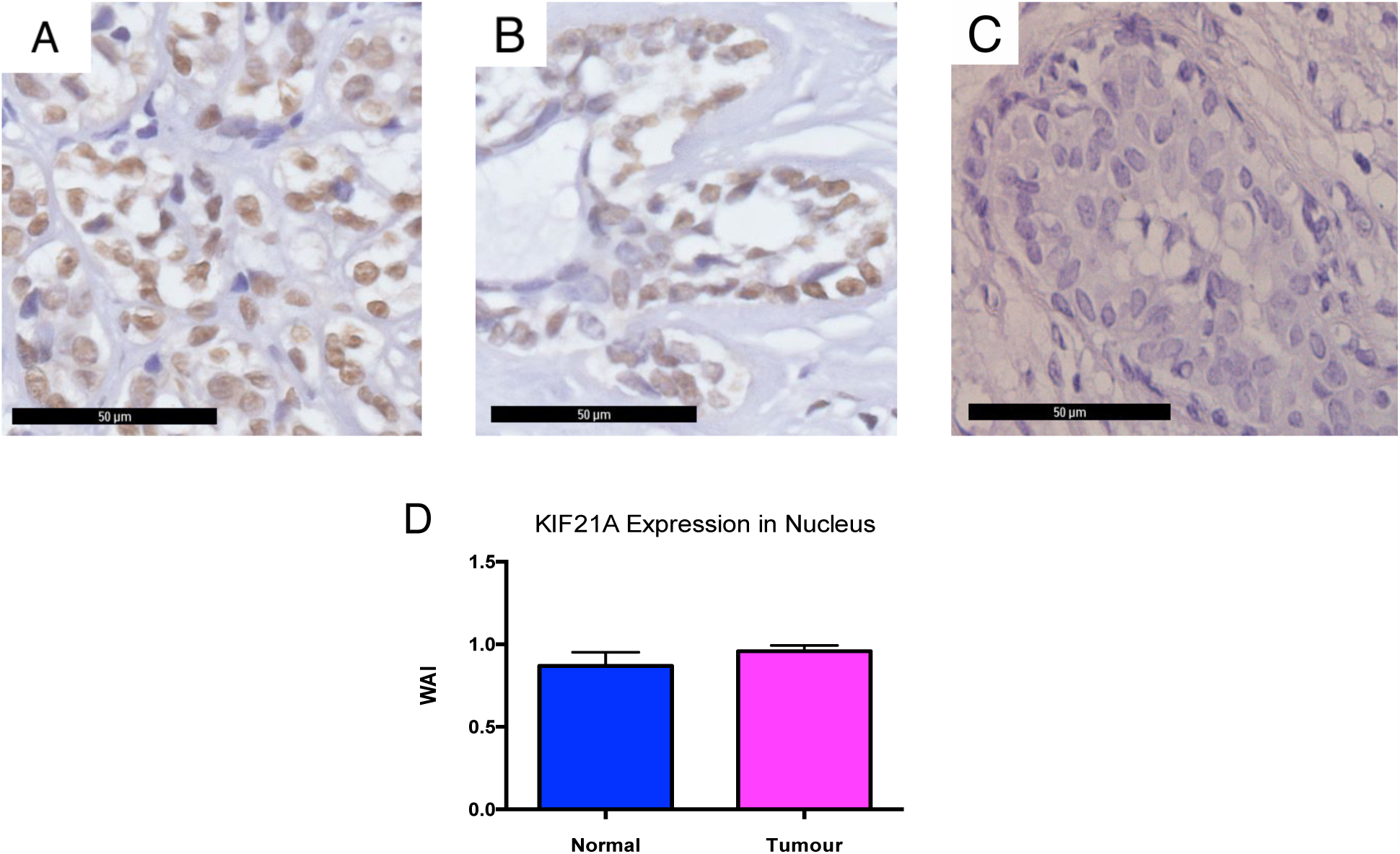
(A,B,C) Representative photomicrographs of KIF21A immunoreactivity in breast IDC tissue. Staining was predominately localised to the epithelial nucleus of (A) tumour cells and (B) adjacent normal tissue, while cytoplasmic staining was rare. (C) Negative control slide. Scale bars represent 50μm. All photomicrographs are at 40x magnification. (D) There were no significant differences in KIF21A protein expression between breast IDC tumour tissues and adjacent normal tissues, as measured by IHC. Values are mean WAI ± SEM. n = 287 (263 tumour, 24 normal) (Mann-Whitney test). Abbreviations: WAI: Weighted Average Intensity.

### 3.7 Relationship Between KIF21A Expression and Clinicopathological Parameters in Breast IDC Tissues

Cut-off values above and below the mean WAI (1.0) were determined as “high” and “low” KIF21A expression, respectively. High KIF21A expression correlated with tumour size below 20 mm (*p* < 0.05), and progressively lower histological grades (*p* < 0.01; Table 5). No significant correlations were observed between KIF21A and other parameters.

**Table 5.** Clinicopathological parameters of breast IDC cases correlated against KIF21A expression in epithelial nucleus of tumour cells. Abbreviations: HER2: human epidermal growth factor receptor 2. * p < 0.05, ** p < 0.01 (Fisher’s exact test for nominal parameters, Kendall’s tau-c test for ordinal parameters).

| Clinicopathological Parameter |  | Weighted Average Index (WAI) |  |  | p-value |
| --- | --- | --- | --- | --- | --- |
|  |  | ≤ 1.0 | > 1.0 | Cases with no data |  |
|  |  | Number of valid cases (%) |  |  |  |
| Age | ≤ 53 yrs old | 89 (59.7) | 60 (40.3) | 0 | 0.093 |
|  | > 53 yrs old | 78 (68.4) | 36 (31.6) |  |  |
| Race | Chinese | 141 (64.4) | 78 (35.6) | 0 | 0.499 |
|  | Non-Chinese | 26 (59.1) | 18 (40.9) |  |  |
| Tumour Size | ≤ 20 mm | 49 (54.4) | 41 (45.6) | 5 | 0.030* |
|  | > 20 mm | 115 (68.5) | 53 (31.5) |  |  |
| Lymphovascular Invasion | No | 128 (62.7) | 76 (37.3) | 1 | 0.759 |
|  | Yes | 38 (65.5) | 20 (34.5) |  |  |
| Lymph Node Involvement | No | 63 (63.6) | 36 (36.4) | 27 | 0.577 |
|  | Yes | 93 (67.9) | 44 (32.1) |  |  |
| Histological Grade | Grade 1 | 19 (47.5) | 21 (52.5) | 7 | 0.002** |
|  | Grade 2 | 63 (61.8) | 39 (38.2) |  |  |
|  | Grade 3 | 84 (73.7) | 30 (26.3) |  |  |
| Oestrogen Receptor | Negative | 56 (64.4) | 31 (35.6) | 4 | 0.892 |
|  | Positive | 108 (62.8) | 64 (37.2) |  |  |
| Progesterone Receptor | Negative | 83 (65.9) | 43 (34.1) | 4 | 0.440 |
|  | Positive | 81 (60.9) | 52 (39.1) |  |  |
| HER2 | Negative | 102 (64.6) | 56 (35.4) | 44 | 0.166 |
|  | Positive | 33 (54.1) | 28 (45.9) |  |  |

ER: oestrogen receptor; PR: progesterone receptor; HER2: human epidermal growth factor receptor 2.

### 3.8 Univariate (Kaplan-Meier) analysis of survival and KIF21A expression

The three timelines investigated were overall survival (OS), survival after recurrence (SAR), and disease-free survival (DFS). Survival data is cause-specific and was available for all cases, with a follow-up period ranging from 0 months to 156 months. It should be noted that no patients survived after breast cancer recurrence for the entire follow-up period (Table 6). Surprisingly, Kaplan-Meier analysis showed patients with high nuclear KIF21A expression were more likely to have better overall survival (*p* < 0.05; Figure 6A) and survival after recurrence (*p* < 0.05; Figure 6B). However, nuclear KIF21A expression was not identified as a significant predictor of breast cancer recurrence, as measured by DFS (Figure 6C).

**Figure 6.**
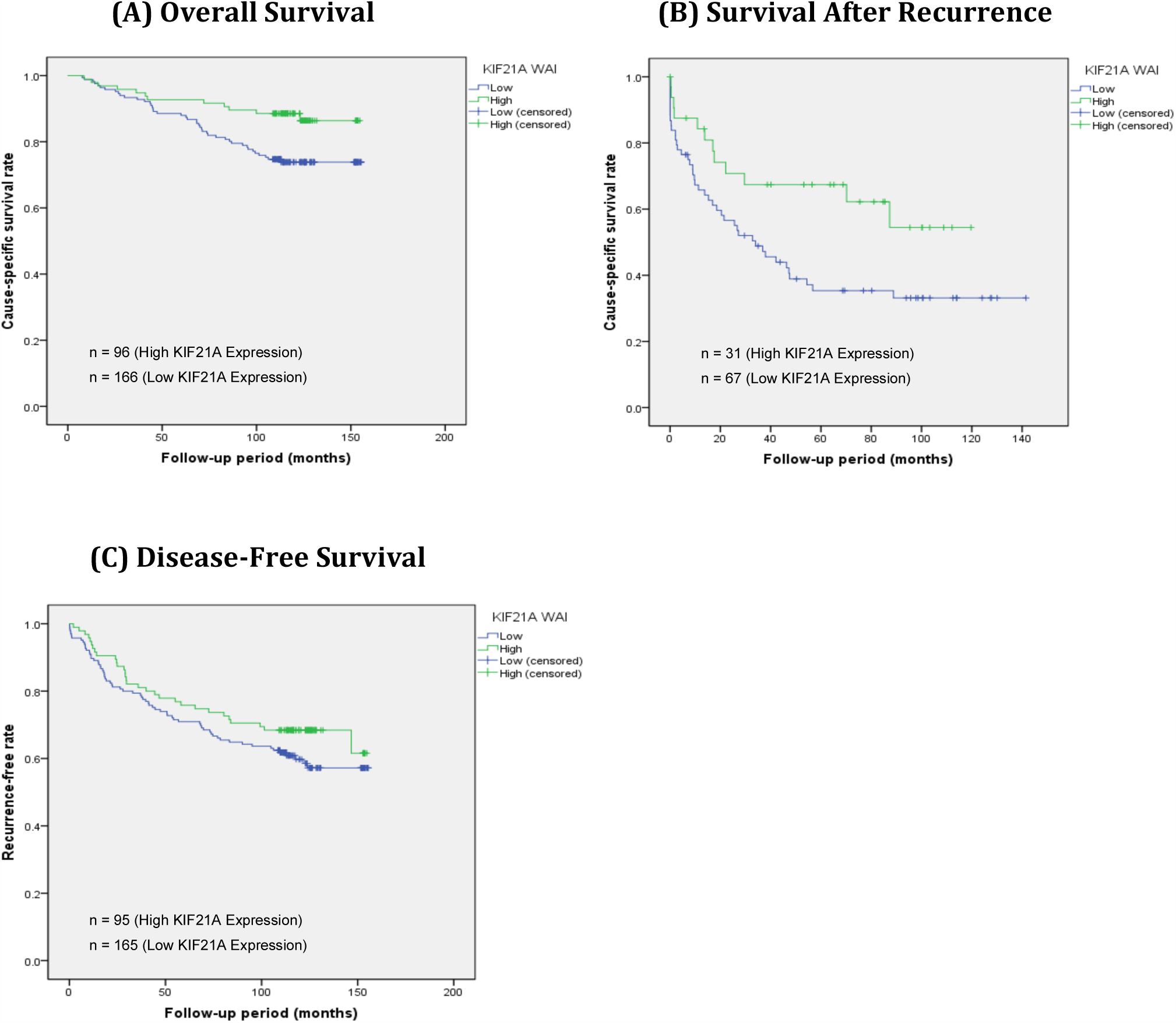
Kaplan-Meier curves for overall cause-specific survival rate (A), cause-specific survival rate after recurrence (B), and disease-free survival (C). KIF21A protein levels in the epithelial nuclei of tumour cells showed prognostic roles in survival (A,B), but not recurrence (C). Patients with low KIF21A expression in breast cancer tissue had significantly shorter survival than those with high KIF21A expression.

**Table 6.** Data summary of nuclear KIF21A epithelial staining, cause-specific mortality (OS and SAR) and recurrence (DFS) in breast IDC patients over 156-month follow-up period. An event is defined as either cause-specific death (for OS and SAR timelines) or recurrence (DFS timeline), while censored cases represent patients who either did not experience an event during the entire follow-up period, or who withdrew from the study during the follow-up period without experiencing an event. * *Log*-*rank p* < 0.05.

| Timeline | KIF21A WAI | Total | No. of events | Censored | | Log-rank $p$ (Mantel-Cox) |
| --- | --- | --- | --- | --- | --- | --- |
|  |  |  |  | No. | Percent |  |
| Overall Survival | ≤ 1.0 | 166 | 43 | 123 | 74.1% | **0.012\*** |
|  | >1.0 | 96 | 12 | 84 | 87.5% |  |
|  | Overall | 262 | 55 | 207 | 79.0% |  |
| Survival After Recurrence | ≤ 1.0 | 67 | 43 | 25 | 37.3% | **0.025\*** |
|  | >1.0 | 31 | 12 | 21 | 67.7% |  |
|  | Overall | 98 | 55 | 46 | 46.9% |  |
| Disease-free Survival | ≤ 1.0 | 165 | 67 | 98 | 59.4% | 0.187 |
|  | >1.0 | 95 | 31 | 64 | 67.4% |  |
|  | Overall | 260 | 98 | 162 | 62.3% |  |

### 3.9 Multivariate (Cox proportional hazards regression) analysis of survival and KIF21A expression

Variables that achieved statistical significance in univariate analyses (*p* < 0.05) were subsequently entered into a multivariate analysis using Cox proportional hazards model via the backward stepwise regression method (Model A; Table 7). In addition, to examine the prognostic value of KIF21A in greater detail, the analysis included a multivariate Cox proportional hazards model that included *all* clinicopathological parameters via the enter method (Model B; Table 7). This included patients’ age, race, tumour size, histological grade, lymph node involvement, lymphovascular invasion, and ER, PR and HER2 statuses. However, only those which showed statistical significance (*p* < 0.05) are shown in Table 7. Model A showed KIF21A acts independently to predict SAR (*p* < 0.05), but significantly correlates with tumour size in predicting OS (*p* < 0.05). Model B predicted KIF21A as an independent prognostic factor for both OS (*p* < 0.05) and SAR (*p* < 0.05). However, both models showed KIF21A expression was not a prognostic factor for recurrence, as measured by DFS.

**Table 7.** Univariate (Kaplan-Meier) and multivariate (Cox proportional hazards regression) analysis of cause-specific survival in breast IDC patients. Abbreviations: HR: hazard ratio; CI: confidence interval; ER: oestrogen receptor. ^a^ Reference groups (HR = 1) were used as the denominator in hazard ratio calculations; ^b^ Histological grades 1 and 2 were binned into “low” grade, while “high” is classified as grade 3; ^c^ Kaplan-Meier univariate analysis performed using the log-rank (Mantel-Cox) test; ^d^ Model A: Cox proportional hazards model including only significant (*p* < 0.05) variables identified from the Kaplan-Meier univariate analysis; ^e^ Model B: Cox proportional hazards model including all variables. * p < 0.05, ** p < 0.01.

| Timeline | Parameter | Reference Group<sup>a</sup><br>(HR = 1) | Univariate Analysis<sup>c</sup> |  |  | Multivariate Analyses |  |  |  |  |  |
| --- | --- | --- | --- | --- | --- | --- | --- | --- | --- | --- | --- |
|  |  |  |  |  |  | Model A<sup>d</sup> |  |  | Model B<sup>e</sup> |  |  |
|  |  |  | HR | 95% CI | p value | HR | 95% CI | p value | HR | 95% CI | p value |
| Overall survival | Tumour Size | ≤ 20 mm | 2.91 | 1.42 – 5.96 | **0.002\*\*** | 2.55 | 1.24 – 5.25 | **0.011\*** | 2.20 | 0.88 – 5.50 | 0.090 |
|  | Histological Grade | Low<sup>b</sup> | 1.67 | 1.10 – 2.54 | **0.012\*** | 1.42 | 0.91 – 2.21 | 0.121 | 0.99 | 0.57 – 1.72 | 0.968 |
|  | KIF21A WAI | Low | 0.45 | 0.24 – 0.85 | **0.012\*** | 0.39 | 0.19 – 0.81 | **0.012\*** | 0.34 | 0.14 – 0.82 | **0.017\*** |
| Survival after recurrence | Tumour Size | ≤ 20 mm | 2.24 | 1.09 – 4.590 | **0.023\*** | 2.03 | 0.99 – 4.19 | 0.055 | 1.30 | 0.49 – 3.45 | 0.604 |
|  | KIF21A WAI | Low | 0.49 | 0.26 – 0.93 | **0.025\*** | 0.52 | 0.27 – 1.01 | **0.049\*** | 0.37 | 0.13 – 1.04 | **0.048\*** |
| Disease-free survival | Tumour Size | ≤ 20 mm | 1.59 | 1.01 – 2.50 | **0.042\*** | 1.21 | 0.72 – 2.03 | 0.477 | 1.29 | 0.77 – 2.14 | 0.335 |
|  | Lymph Node Involvement | No | 1.83 | 1.16 – 2.88 | **0.008\*\*** | 1.44 | 1.18 – 1.75 | **0.001\*\*** | 1.87 | 1.17 – 2.98 | **0.009\*\*** |
|  | Histological Grade | Low<sup>b</sup> | 1.53 | 1.13 – 2.07 | **0.021\*** | 1.22 | 0.86 – 1.74 | 0.261 | 1.30 | 0.92 – 1.83 | 0.142 |
|  | ER | Negative | 0.50 | 0.33 – 0.74 | **0.001\*\*** | 0.47 | 0.30 – 0.73 | **0.001\*\*** | 0.56 | 0.29 – 1.07 | **0.001\*\*** |
|  | KIF21A WAI | Low | 0.75 | 0.49 – 1.15 | 0.187 | - | - | - | 0.47 | 0.30 – 0.73 | 0.080 |

## 4 DISCUSSION

### 4.1 KIF21A Mediates Breast Cancer Metastasis *In Vitro*

This study identifies KIF21A as a protein whose depletion consistently caused a statistically and biologically significant reduction in breast cancer cell migration and invasion *in vitro* (Figures 2 and 3). To our knowledge, this is the first study to identify the role of KIF21A in cancer cell metastasis – an important finding in the assessment of KIF21A as a potential therapeutic target. Invasion of malignant cells into the surrounding ECM and migration of invaded cells towards the bloodstream are pivotal early steps in metastasis, suggesting that KIF21A is an important mediator of early carcinogenesis. The observed migratory influence of KIF21A was enhanced in MDA-MB-231 cells, which are known to be highly metastatic, while higher silencing efficiency using siKIF21A-2 in MCF-7 intensified the reduction in migration. Furthermore, although a more optimal silencing efficiency would be preferred to observe the full effects of KIF21A knockdown, there was still a significant reduction in both invasion and migration. Whether complete knockdown would have accentuated this observation remains unknown. Nonetheless, these cumulative findings suggest that KIF21A may be a key component of breast cancer cell migratory and invasive pathways. This could be explained by its interaction with other molecules and their known mechanisms (see Section 4.3 below).

However, these observations surprisingly contradicted the *in vivo* findings of KIF21A expression in human breast cancer tissue (discussed in-detail in Section 4.2), meaning that KIF21A’s role in cancer metastasis is largely inconclusive. Despite this, *in vitro* functional assays do serve an important purpose. Although they are only representative of complex *in vivo* conditions, they provide a critical understanding about the influence of genes on distinct metastatic components. In this case, KIF21A was shown to influence the distinct metastatic components of migration and invasion – a phenomenon supported by studies of other kinesin family members. KIF3A and KIF3B, for example, have been shown to interact in a complex that transports proteins essential for cancer cell migration [23], while KIF11 has been found to respond to directional cues during chemotaxis to govern the direction of MDA-MB-231 cell migration [24]. Other studies identify KIF14 [25], KIF2A [26], and KIF18A [27] as mediators of breast cancer migration *in vitro* via a range of pathways. Given the discrepant results of the current study, migration and invasion assays should be extended to *in vivo* models to elucidate KIF21A’s precise role in these processes.

Following migration, cancer cells must adhere to the ECM of a new site where they can then rapidly proliferate. This study investigated the role of KIF21A in this process. KIF21A knockdown by 40-70% did not induce any changes in cell proliferation for both breast cancer cell lines (Figure 4), and thus KIF21A is not implicated as a player in breast cell proliferation. Although it is difficult to identify studies reporting similarly negative results for other kinesin family members, the present study shows KIF21A varies from previous reports of KIFs acting as key mediators of proliferation in breast cancer [26,28].

KIF21A was also tested for cell adhesion to collagen I and fibronectin fibers – both protein components of the ECM – yet no changes were observed following silencing in MCF-7 for both fibers, and in MDA-MB-231 for collagen I fibers (Figure 4). However, reduced adhesion to fibronectin did occur in MDA-MB-231 cells, suggesting KIF21A could facilitate breast cancer cell establishment in foreign sites, particularly for highly invasive cells. The fundamental genetic differences between MCF-7 and MDA-MB-231 cells may have contributed to differences with their adhesion assays. Further analysis is thus required to investigate the functional role of KIF21A in breast cancer cell adhesion, which could be done via overexpression studies and other adhesion assays involving laminin, gelatin and collagen II fibers. More broadly, there is no particular commonality in the adhesive capabilities of other kinesins, with many being shown to either positively or negatively regulate adhesion in various cell lines [25,29,30]. Further studies in KIF21A are therefore encouraged.

### 4.2 Disparity Between *In Vitro* and *In Vivo* Results

The present study suggested KIF21A exerts oncogenic activity *in vitro,* which resonates with our hypothesis and previous studies that show KIF21A facilitates the survival of breast cancer cell lines [22]. However, conflicting results were obtained *in vivo.* Immunohistochemical analysis suggested KIF21A exerts tumour-suppressive functions, rather than oncogenic activity. Although it is unclear why different results were obtained for the same gene, there are several explanations for this phenomenon: (1) ethnic differences. Various studies illustrate that the mutation status of numerous genes contributing to breast cancer is different between Asian and Caucasian populations. This study was predominately performed in Asian individuals, while MCF-7 cells and MDA-MB-231 cells were isolated from Caucasian women. Ethnicity may therefore be a contributing factor to any discrepancies, and *in vivo* studies of KIF21A in Caucasian populations are encouraged. (2) One molecule may have dual or multiple physiological functions. For example, both transforming growth factor beta (TGF-beta) and signal transducer and activator of transcription 3 (STAT3) have oncogenic or tumour-suppressive roles depending on different conditions including the mutational background of the tumor [31–33]. Whether KIF21A has a dual effect remains unknown, however similarly conflicting studies have been observed in other kinesins, particularly KIF14, which has been shown to both promote and suppress tumorigenesis [15,34]. (3) Study limitations. *In vitro* assays are useful in studying the influence of various molecules on distinct metastatic components, but we must not forget the amazingly complex story of the tumour microenvironment. Firstly, despite strict conditions, cell line studies cannot control for the vast array of biological parameters that would otherwise be present *in vivo*. Cell lines studies are therefore better used as tools to delineate the molecular events underpinning carcinogenesis rather than predicting the entire metastatic cascade. Secondly, cell lines are only representative of one individual. Although two breast cancer cell lines were used for support in this study, they cannot match the statistical power of the 263 breast IDC patient samples analysed via IHC. Of course, some patients may have indeed had a poorer prognosis with high KIF21A expression, which would resonate with the *in vitro* findings. However, there was a significant trend towards poorer prognosis correlating with low KIF21A expression. Further *in vitro* analyses are therefore encouraged in more breast cancer cell lines that are subject to both KIF21A silencing and overexpression.

### 4.3 KIF21A Signaling Pathways Leading to Cell Migration and Invasion

Although few studies have investigated KIF21A beyond its role in congenital fibrosis of the extraocular muscles type-1 (CFEOM1), evidence suggests it may interact with other genes
involved in cell migration. KIF21A binds to brefeldin A (BFA)-inhibited guanine nucleotide-exchange factor (BIG1) [35] to maintain the organisation of the Golgi apparatus [36], and also transports KN motif and ankyrin repeat domains 1 (KANK1), influencing its membrane localisation [37]. Subsequent exploration revealed that both BIG1 and KANK1 coimmunoprecipitated with KIF21A and each other, such that they may act as a scaffold in KIF21A-powered intracellular transport [38]. The interaction between all three molecules was shown to affect cell migration; a phenomenon explained by their combined effects on cell polarity. Generation and maintenance of cell polarity is essential for directional migration and results from asymmetric membrane traffic achieved by intracellular transport and the delivery of extra membrane to the cell’s leading edge [39,40]. KIF21A is theorised to act via BIG1 and KANK1 to position the Golgi and microtubule-organising center (MTOC) structures anterior to the nucleus, resulting in a polarity shift that induces front-end-directed cell migration [38]. This interaction would explain the observations seen in breast cancer cell migration in the present study. In support of this theory, previous studies show polarisation of Golgi and MTOC structures to be explicitly disturbed by KANK1 or BIG1 siRNA treatment [38]. However, further investigation is required to identify additional players in KIF21A/BIG1/KANK1 functional interactions and to elucidate the detailed molecular mechanisms that alter cell polarity to influence directional migration.

### 4.4 KIF21A in Breast Cancer Tissues

Although our *in vitro* data strongly supported a tumorigenic role for KIF21A in breast cancer, IHC in breast cancer tissue microarrays was contrary to our hypothesis and showed KIF21A associated with anti-malignant phenotypes and a better prognosis. Both analyses suggest KIF21A is involved in breast cancer pathogenesis and prognosis, however their conflicting outcomes mean the overall results of this study remain largely inconclusive.

Nonetheless, numerous important findings came from the IHC analysis. Firstly, there were no significant differences in KIF21A expression between breast IDC samples and adjacent normal tissues (Figure 5). This adds to inconsistent evidence regarding the role of KIF21A in breast oncogenesis, and is contradictory to previous studies in the kinesin superfamily, which observe upregulation of numerous KIFs in breast cancer [13,15,17]. KIF21A may therefore not be a useful diagnostic biomarker.

In addition, KIF21A expression was shown to negatively correlate with the pro-malignant phenotypes of large tumour size (≥ 20 mm) and high histological grade (Table 5). Large tumour size and high histological grade classically predict poorer prognosis in breast cancer [41], so KIF21A may exert a tumour-suppressive role during breast carcinogenesis. However, the mechanism via which it performs this role was not clear from the IHC analysis, as KIF21A displayed no relationships with lymphovascular invasion, lymph node involvement, or various receptors including ER, PR, and HER2. This suggests KIF21A may not regulate distant metastasis to lymph nodes and blood vessels (in contradiction to the *in vitro* observations), and it may not be involved in the mechanisms through which oestrogen, progesterone and human epidermal growth factor increase breast cell proliferation. This differs from previous observations of concerted KIF21A upregulation upon the introduction of exogenous oestrogen *in vitro* [42]. One explanation for KIF21A-induced tumour suppression may therefore be its role in mitosis (see Section 4.5 below).

Survival analyses were also performed, and revealed high KIF21A expression predicted better OS and SAR, but had no significant associations with DFS (Figure 6). KIF21A may therefore act as a tumour suppressor, and may be used as a predictor of breast cancer survival, but not recurrence. To identify whether KIF21A predicted survival independently of other clinicopathological parameters, multivariate analyses were also performed (Table 7). Although univariate analysis suggested KIF21A co-predicted OS and SAR with tumour size and histological grade, Cox proportional hazards models revealed KIF21A acted independently. KIF21A may therefore be an independent prognostic biomarker for better breast cancer survival. In this study cohort, lymphovascular invasion, lymph node involvement, and hormonal and growth factor receptor statuses had no correlations with SAR and OS – a rare finding (although some of those variables did correlate with recurrence). Nonetheless, this observation adds to evidence that KIF21A functions independently of lymphovascular invasive pathways, oestrogen, progesterone, and human epidermal growth factor to implement its tumour-suppressive activity.

The possible role of KIF21A as a tumour suppressor and predictor of better breast cancer prognosis differs from that of most other kinesin family members. KIFs 2A [43], 2C [17], 3C [44], and 26B [45] have all been identified as predictors of worse breast cancer prognosis. Intriguingly however, a select few kinesins have been observed as tumour-suppressor genes consistent with the results of the current study, albeit in other cancer types. Overexpression of both KIF4 [46] and KIF14 [34] has been shown to reduce metastatic phenotypes *in vitro* and predict better survival outcomes in human gastric carcinoma and lung adenocarcinoma, respectively. In both cases, it was theorised that they carried out their tumour-suppressive functions through their roles in mitosis – a phenomenon that could explain similar observations in this study (see below).

### 4.5 Mitotic Misregulation as a Functional Explanation for KIF21A-mediated Tumorigenesis

Errors at any point during the cell cycle can be catastrophic and in humans can lead to cancer [47]. Chromosome mis-segregation during mitosis, for example, results in abnormal cytokinesis and aneuploidy [48], which is cleared in normal cells through apoptosis. However, tumour cells show higher rates of aneuploidy, which is often associated with poor clinical outcomes [49–51]. Interestingly, misregulation of various mitotic kinesins has been shown to result in aneuploidy, because unbalanced movement of kinesins can cause excessive spindle separation, premature sister chromatid detachment, overshooting before anaphase, and eventually unequal distribution of DNA [8–11]. The aneuploid daughter cells could, theoretically, display any possible tumorigenic phenotype and a plethora of metastatic characteristics, including aberrant migration and invasion. Via this mechanism, KIF21A depletion could result in aneuploidy to facilitate tumour formation, which explains the *in vivo* findings of this study. High KIF21A expression may therefore suppress carcinogenesis if KIF21A is implicated in mitosis.

### 4.6 Future Work

The results of this study were largely inconclusive due to discrepancies between *in vitro* and clinical data. To further elucidate KIF21A’s role in breast cancer, other functional analyses should therefore be performed (e.g. apoptosis and drug resistance assays) in more breast cancer cell lines that have been subjected to KIF21A knockdown, overexpression and the introduction of exogenous KIF21A. This would solidify knowledge about KIF21A’s role in breast cancer *in vitro*, however there still remains little physiological evidence on the role of KIF21A *in vivo*. IHC analysis of an Asian population in this study should therefore be extended to Caucasian cohorts. Furthermore, given many known KIF proteins occur in mice [4], they would be an ideal animal model to knockout genes and explore physiological effects in tumour xenografts. KIF21A’s biological pathways are also poorly understood, and could be discovered through microarray analysis of KIF21A-silenced cells. Any upstream or downstream targets of KIF21A that are known breast cancer oncogenes would provoke interesting follow-up studies, and have the potential to be targeted alongside KIF21A in combinative therapy.

Because of the lack of primary data, KIF21A and many other kinesins are yet to be considered therapeutic targets or prognostic predictors, despite common consensus that kinesins play a vital role in breast carcinogenesis [8]. If mounting evidence supports either a tumour-suppressive or oncogenic role for KIF21A in the metastatic cascade, its pathways could be modulated through targeted chemotherapeutic strategies. Future studies should therefore attempt to uncover the structure of KIF21A’s ATP-, microtubule and enzymebinding sequences, which could lead to the development of KIF21A inhibitors and enhancers. Identifying binding partners has enormous therapeutic potential, because drugs that mimic those partners could bind to allosteric pockets to either promote or suppress KIF21A and its effects on carcinogenesis.

### 4.7 Conclusions

In summary, this study illustrates the potential involvement of KIF21A in breast cancer pathogenesis and progression. However, the conflicting outcomes of *in vitro* and *in vivo* data means no definitive conclusions can be drawn about KIF21A’s role in breast cancer. This may be due to ethnic differences, the possible multi-functionality of KIF21A, or even study limitations, and warrants further investigation into the influence of KIF21A *in vivo,* the molecules it interacts with, and its potential as a prognostic biomarker. Such knowledge could lead to the development of novel chemotherapeutic strategies to mediate its function and enhance prognostic outcomes.

## REFERENCES

1. Ferlay J, Soerjomataram I, Ervik M, Dikshit R, Eser S, Mathers C, et al. GLOBOCAN 2012 v1.0, Cancer Incidence and Mortality Worldwide: IARC CancerBase No. 11 [Internet]. Lyon, France: International Agency for Research on Cancer; 2013. Available from: http://globocan.iarc.fr, accessed on 01/09/2015.

2. Liu X, Gong H, Huang K. Oncogenic role of kinesin proteins and targeting kinesin therapy. Cancer Sci. 2013 Jun;104(6):651–6.

3. Vale RD, Reese TS, Sheetz MP. Identification of a novel force-generating protein, kinesin, involved in microtubule-based motility. Cell. 1985 Aug;42(1):39–50.

4. Miki H, Setou M, Kaneshiro, Hirokawa. All kinesin superfamily protein, KIF, genes in mouse and human. Proc Natl Acad Sci USA. 2001 Jun 19;98(13): 7004–11.

5. Lawrence CJ, Dawe RK, Christie KR, Cleveland DW, Dawson SC, Endow S, et al. A standardized kinesin nomenclature. J Cell Biol. 2004 Oct 11;167(1):19–22.

6. Diefenbach RJ, Mackay JP, Armati PJ, Cunningham AL. The C-terminal region of the stalk domain of ubiquitous human kinesin heavy chain contains the binding site for kinesin light chain. Biochemistry. 1998;37(1):16663–70.

7. Goldstein LS, Philp AV. The road less traveled: emerging principles of kinesin motor utilization. Annu Rev Cell Dev Biol. 1999;15(2): 141–83.

8. Castillo A, Morse HC, Godfrey VL, Naeem R, Justice MJ. Overexpression of Eg5 causes genomic instability and tumour formation in mice. Cancer Res. 2007;67(21):10138–47.

9. Wordeman L. How kinesin motor proteins drive mitotic spindle function: lessons from molecular assays. Semin Cell Dev Biol 2010;21(3):260–8.

10. Carleton M, Mao M, Biery M, Warrener P, Kim S, Buser C, et al. RNA interference-mediated silencing of mitotic kinesin KIF14 disrupts cell cycle progression and induces cytokinesis failure. Mol Cell Biol. 2006 May;26(10):3853–63.

11. Molina I, Baars S, Brill JA, Hales KG, Fuller MT, Ripoll P. A chromatin-associated kinesin-related protein required for normal mitotic chromosome segregation in Drosophila. J Cell Biol. 1997 Dec 15;139(6):1361–71.

12. Oki E, Hisamatsu Y, Ando K, Saeki H, Kakeji Y, Maehara Y. Clinical aspect and molecular mechanism of DNA aneuploidy in gastric cancers. J. Gastroenterol 2012;47(4):351–8.

13. Wu G, Zhou L, Khidr L, Guo XE, Kim W, Lee YM, et al. A novel role of the chromokinesin KIF4A in DNA damage response. Cell Cycle. 2008;7(13):2013–2020.

14. Mazumdar M, Lee JH, Sengupta K, Ried T, Rane S, Misteli T. Tumor formation via loss of a molecular motor protein. Curr Biol. 2006 Aug 8;16(15):1559–64.

15. Corson TW. Gallie BL. KIF14 mRNA expression is a predictor of grade and outcome in breast cancer. Int J Cancer. 2006 Sep 1;119(5):1088–94.

16. Sanhaji M, Friel CT, Wordeman L, Louwen F, Yuan J. Mitotic centromere-associated kinesin (MCAK): a potential cancer drug target. Oncotarget. 2011 Dec;2(12):935–47.

17. Nishidate T, Katagiri T, Lin ML, Mano Y, Miki Y, Kasumi F, et al. Genome-wide geneexpression profiles of breast-cancer cells purified with laser microbeam microdissection: identification of genes associated with progression and metastasis. Int J Oncol. 2004 Oct;25(4):797–819.

18. Shimo A, Tanikawa C, Nishidate T, Lin ML, Matsudi K, Park JH, et al. Involvement of kinesin family member 2C/mitotic centromere-associated kinesin overexpression in mammary carcinogenesis. Cancer Sci. 2008 Jan;99(1):62–70.

19. De S Cipriano R, Cipriano R, Jackson MW, Stark GR. Overexpression of kinesins mediates docetaxel resistance in breast cancer cells. Cancer Res. 2009 Oct;69(20):8035–42.

20. Tan MH, De S, Bebek G, Orloff MS, Wesolowski R, Downs-Kelly E, et al. Specific kinesin expression profiles associated with taxane resistance in basal-like breast cancer. Breast Cancer Res Treat. 2012 Feb;131(3):849–58.

21. Ganguly A, Yang H, Cabral F. Overexpression of mitotic centromere-associated kinesin stimulates microtubule detachment and confers resistance to paclitaxel. Mol Cancer Ther. 2011 Jun;10(6):929–37.

22. Groth-Pedersen L, Aits S, Corcelle-Termeau E, Petersen NH, Nylandsted J, Jäättelä M. Identification of cytoskeleton-associated proteins essential for lysosomal stability and survival of human cancer cells. PLoS One. 2012;7(10):e45381.

23. Jimbo T, Kawasaki Y, Koyama R, Sato R, Takada S, Haraguchi K, et al. Identification of a link between the tumour suppressor APC and the kinesin superfamily. Nat Cell Biol. 2002 Apr;4(4):323–7.

24. Wang F, Lin SL. Knockdown of kinesin KIF11 abrogates directed migration in response to epidermal growth factor-mediated chemotaxis. Biochem Biophys Res Commun. 2014 Sep;452(3):642–8.

25. Ahmed SM, Theriault BL, Uppalapati M, Chiu CW, Gallie BL, Sidhu SS, et al. KIF14 negatively regulates Rap1a-Radil signaling during breast cancer progression. J Cell Biol. 2012 Dec 10;199(6):951–67.

26. Wang J, Ma S, Ma R, Qu X, Liu W, Ly C, et al. KIF2A silencing inhibits the proliferation and migration of breast cancer cells and correlates with unfavorable prognosis in breast cancer. BMC Cancer. 2014 Jun 21;14:461.

27. Zhang C, Zhu C, Chen H, Li L, Guo L, Jiang W, et al. Kif18A is involved in human breast carcinogenesis. Carcinogenesis. 2010 Sep;31(9):1676–84.

28. Yu Y, Wang XY, Sun L, Wang YL, Wan YF, Li XQ, et al. Inhibition of KIF22 suppresses cancer cell proliferation by delaying mitotic exit through upregulating CDC25C expression. Carcinogenesis. 2014 Jun;35(6):1416–25.

29. Li G, Luna C, Qiu J, Epstein DL, Gonzalez P. Targeting of Integrin β1 and Kinesin 2α by MicroRNA 183. J Biol Chem. 2010 Feb 19; 285(8): 5461–5471.

30. Yoon JR, Whipple RA, Balzer EM, Cho EH, Matrone MA, Peckham M, et al. Local anesthetics inhibit kinesin motility and microtentacle protrusions in human epithelial and breast tumor cells. Breast Cancer Res Treat. 2011 Oct;129(3):691–701.

31. Wakefield LM, Roberts AB. TGF-beta signaling: positive and negative effects on tumorigenesis. Curr Opin Genet Dev. 2002 Feb;12(1):22–9.

32. Roberts AB, Wakefield LM. The two faces of transforming growth factor beta in carcinogenesis. Proc Natl Acad Sci USA. 2003 Jul 22;100(15):8621–3.

33. de la Iglesia N, Konopka G, Puram SV, Chan JA, Bachoo RM, You MJ, et al. Identification of a PTEN-regulated STAT3 brain tumor suppressor pathway. Genes Dev. 2008 Feb 15;22(4):449–62.

34. Hung PF, Hong TM, Hsu YC, Chen HY, Chang YL, Wu CT, et al. The motor protein KIF14 inhibits tumor growth and cancer metastasis in lung adenocarcinoma. PLoS One. 2013;8(4):e61664.

35. Shen X, Meza-Carmen V, Puxeddu E, Wang G, Moss J, Vaughan M. Interaction of brefeldin Ainhibited guanine nucleotide-exchange protein (BIG) 1 and kinesin motor protein KIF21A. Proc Natl Acad Sci USA. 2008 Dec 2; 105(48):18788–93.

36. Boal F, Stephens DJ. Specific functions of BIG1 and BIG2 in endomembrane organization. PLoS One. 2010 Mar 25; 5(3):e9898.

37. Kakinuma N, Kiyama R. A major mutation of KIF21A associated with congenital fibrosis of the extraocular muscles type 1 (CFEOM1) enhances translocation of Kank1 to the membrane. Biochem Biophys Res Commun. 2009 Sep 4; 386(4):639–44.

38. Li CC, Kuo JC, Waterman CM, Kiyama R, Moss J, Vaughan M. Effects of brefeldin A-inhibited guanine nucleotide-exchange (BIG) 1 and KANK1 proteins on cell polarity and directed migration during wound healing. Proc Natl Acad Sci U S A. 2011 Nov 29;108(48):19228–33.

39. Etienne-Manneville S. Cdc42--the centre of polarity. J Cell Sci. 2004 Mar 15;117(8):1291–300.

40. Etienne-Manneville S, Hall A. Cell polarity: Par6, aPKC and cytoskeletal crosstalk. Curr Opin Cell Biol. 2003 Feb;15(1):67–72.

41. Narod SA. Tumour size predicts long-term survival among women with lymph node-positive breast cancer. Curr Oncol. 2012 Oct; 19(5):249–253.

42. Zou JX, Duan Z, Wang J, Sokolov A, Xu J, Chen CZ, Li JJ, Chen HW. Kinesin family deregulation coordinated by bromodomain protein ANCCA and histone methyltransferase MLL for breast cancer cell growth, survival, and tamoxifen resistance. Mol Cancer Res. 2014 Apr;12(4):539–49.

43. Wang J, Ma S, Ma R, Qu X, Liu W, Ly C, et al. KIF2A silencing inhibits the proliferation and migration of breast cancer cells and correlates with unfavorable prognosis in breast cancer. BMC Cancer. 2014 Jun 21;14:461.

44. Wang C, Wang C, Wei Z, Li Y, Wang W, Li X, et al. Suppression of motor protein KIF3C expression inhibits tumor growth and metastasis in breast cancer by inhibiting TGF-β signaling. Cancer Lett. 2015 Nov 1;368(1):105–14.

45. Wang Q, Zhao ZB, Wang G, Hui Z, Ming-Hua W, Jun-Feng P, et al. High expression of KIF26B in breast cancer associates with poor prognosis. PLoS One. 2013;8(4):e61640.

46. Gao J, Sai N, Wang C, Sheng X, Shao Q, Zhou C, et al. Overexpression of chromokinesin KIF4 inhibits proliferation of human gastric carcinoma cells both in vitro and in vivo. Tumour Biol. 2011 Feb;32(1):53–61.

47. Massague J. G1 cell cycle control and cancer. Nature. 2004;432:298–306.

48. Bakhoum SF, Compton DA. Chromosomal instability and cancer: a complex relationship with therapeutic potential. J Clin Invest. 2012 Apr;122(4):1138–43.

49. Chan JY. A clinical overview of centrosome amplification in human cancers. Int J Biol Sci. 2011;7(8):1122–44.

50. McGranahan N, Burrell RA, Endesfelder D, Novelli MR, Swanton C. Cancer chromosomal instability: therapeutic and diagnostic challenges. EMBO Rep. 2012 Jun 1;13(6):528–38.

51. Pfau SJ, Amon A. Chromosomal instability and aneuploidy in cancer: from yeast to man. Rep. 2012 Jun 1;13(6):515–27.

